# A fast variational algorithm to detect the clonal copy number substructure of tumors from single-cell data

**DOI:** 10.1101/2021.11.20.469390

**Authors:** Antonio De Falco, Francesca Caruso, Xiao-Dong Su, Antonio Iavarone, Michele Ceccarelli

## Abstract

Here we report Single CEll Variational ANeuploidy analysis (SCEVAN), a fast variational algorithm for the deconvolution of the clonal substructure of tumors from single cell data. It uses a multichannel segmentation algorithm exploiting the assumption that all the cells in a given copy number clone share the same breakpoints. Thus, the smoothed expression profile of every individual cell constitutes part of the evidence of the copy number profile in each subclone. SCEVAN can automatically and accurately discriminate between malignant and non-malignant cells, resulting in a practical framework to analyze tumors and their microenvironment. We apply SCEVAN to several datasets encompassing 106 samples and 93,322 cells from different tumors types and technologies. We demonstrate its application to characterize the intratumor heterogeneity and geographic evolution of malignant brain tumors.

## Introduction

Understanding intratumor heterogeneity and the interactions between tumor cells and the immune system is the critical step to explain treatment failure and plays a crucial role in the study of tumor growth and evolution^1, 2^. Single-cell RNA sequencing (scRNA-seq) is the reference technology to characterize tumor heterogeneity and the composition of the tumor microenvironment at high resolution^3^. In addition, this technology was successfully used to identify multiple transcriptional programs activated in a single tumor^4, 5^ and to prioritize of key regulators of tumor-host interaction^6^. To study the complexity of lineage identity, differentiation, and proliferation of tumor cells and the impact of stromal and immune components, a large number of unsorted cells from tumor biopsies are subject to whole transcriptomics profiling and then classified as malignant cells, stromal cells, and immune cells, and further stratified into different compartments according either expression of specific markers^6^, and the orchestrated activation of pathways^5^. The distinction of malignant from non-malignant cells is a critical step in the follow-up analysis of scRNA-seq tumor datasets. The basic idea to solve such a problem relies on estimating common copy number alterations that characterize transformed cells. The copy number profiles are obtained by considering the gene expression profiles of each cell as a function of the genomic coordinates. The moving average smoothing of the gene expression function is then clustered in malignant and non-malignant cells. One of the most successful methods based on this approach is the INFERCNV^4^ algorithm. The main drawback is that the clusters of reference cells require manual identification, usually with a combination of approaches^7, 8^. Moreover, INFERCNV and similar methods^4, 9^ are particularly suited for smart-seq data having high coverage and relatively low throughput whereas they exhibit sub-optimal performances on droplet-based methods with very sparse coverage depth and higher throughput^10^. An approach to overcome these limitations is represented by the CopyKAT method^11^ that classifies malignant and non-malignant cells. It was successfully applied to analyze the clonal substructure of three triple-negative breast tumors. However, the classification produced by CopyKAT is severely affected by a wrong identification of normal cells and, similarly to other methods, was not designed to perform a complete automatic identification of the clones, reporting their breakpoints, the specific and shared alteration and a complete e clonal deconvolution.

Here we present Single CEll Variational Aneuploidy aNalysis (SCEVAN), a novel variational algorithm for automatically detecting the clonal copy number substructure of tumors from single-cell data. By overcoming most of the mentioned limitations, our method automatically segregates malignant cells from non-malignant cells. Clusters of malignant cells are then analyzed through an optimization-based joint segmentation algorithm. We exploit the notion that all the cells in a given copy number clone share the same breakpoints with the smoothed expression profile of every individual cell providing support towards the definition of the copy number profile of individual subclones. Therefore, joint segmentation allows the enhancement of systematic biases leading to the emergence of consistent breakpoints. Afterwards, SCEVAN performs a complete downstream analysis to automatically identify tumor subclones, classifying their specific and shared alterations up to a clone phylogeny. The joint segmentation algorithm implemented in SCEVAN is based on a variational framework developed in the field of Computer Vision making use of the Mumford Shah energy model^12^ which has been already successfully applied to detect copy number alterations in matched tumor-normal pairs of high-density comparative genomic hybridizations arrays^13^ and used to detect fusion breakpoints^14^. Moreover, its joint version was developed to identify recurrent copy number alterations in large tumor cohorts^15, 16^. Here we benchmark the output of SCEVAN against state-of-the-art methods and found that SCEVAN exhibits faster and more accurate performance. Finally, we used SCEVAN to characterize the clonal substructure in multiple scRNA-seq glioma and head and neck cancer datasets.

## Results

### SCEVAN Workflow

The workflow of the proposed SCEVAN algorithm (Figure 1) starts from the raw count matrix with genes on rows and cells on columns. The input count matrix is pre-processed by removing cells with a low number of detected transcripts and selecting the most expressed genes. A set of high confident non-malignant cells are identified and used to determine a copy number baseline and to compute the relative matrix removing the baseline (Steps A and B). This matrix undergoes an edge-preserving non-linear diffusion filter assuming a piece-wise smooth function as the underlying model (Step C). The smoothed matrix is then segmented with the joint segmentation algorithm to obtain a copy number matrix (Step D). At this point, SCEVAN discriminates the normal cells from tumor cells as those falling in the cluster containing the highest number of confident normal cells (Step E). Finally, the different subclones are obtained by analyzing the clusters of the tumor cells in the Copy Number Matrix as detailed in the Methods (Step F). In particular, each cluster are segmented independently. The segments are classified and a *p*-value is assigned to each detected alteration. Finally, SCEVAN characterizes truncal, shared and clone-specific alterations comparing different clusters, performing enrichment analysis up to a clone phylogeny.

**Figure 1.**
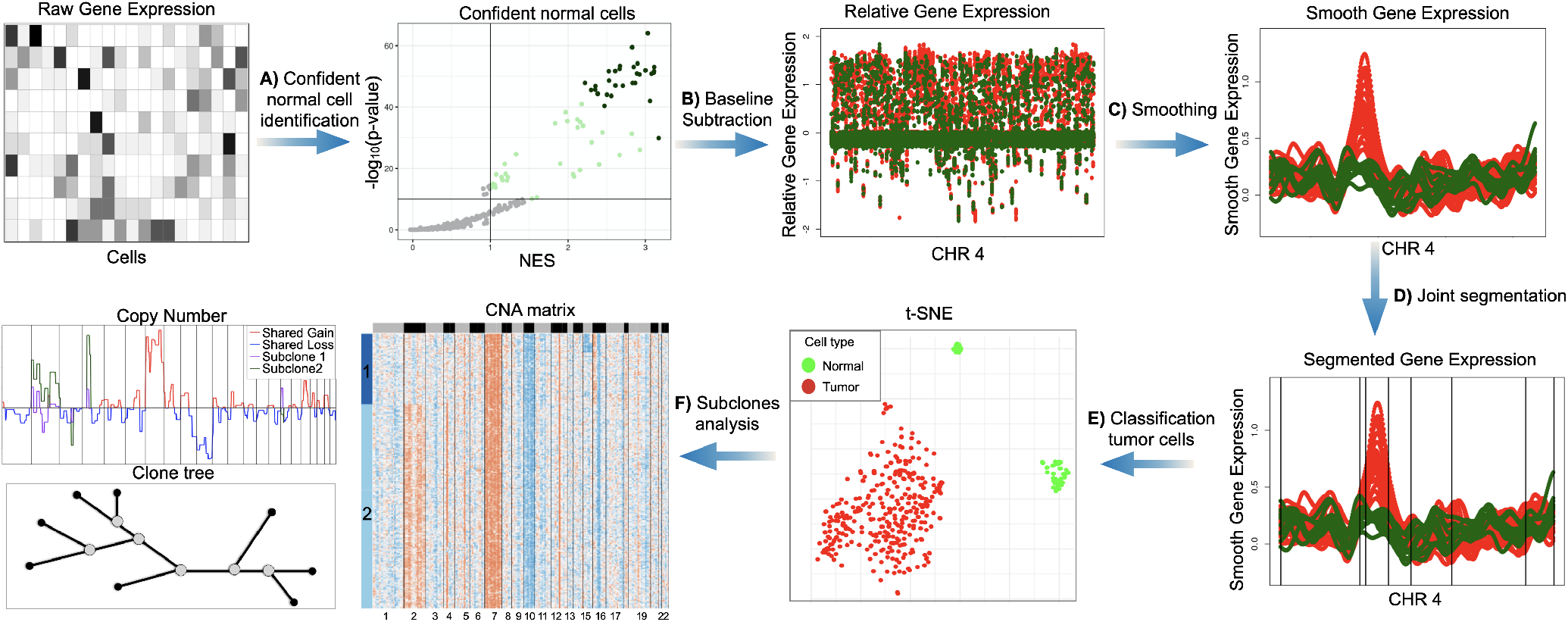
SCEVAN starts from the raw count matrix removing irrelevant genes and cells. **A)** Identification of a small set of highly confident normal cells. **B)** Relative gene expression obtained from removal of the baseline inferred from confident normal cells. **C)** Edge-preserving non-linear diffusion filtering of relative gene expression. **D)** Segmentation with a a variational region growing algorithm. **E)** identification of normal cells as those in the cluster containing the majority of confident normal cells. **F)** Analysis of subclones within tumour cells and characterization of shared and specific alterations.

### Malignant cell classification

To evaluate the accuracy of SCEVAN in classifying malignant from non-malignant cells, we applied out tool to several public datasets^7, 10, 17–19^ of three different cancer type of scRNA-seq data (Glioblastoma (GBM), Head and Neck Squamous Cell Carcinomas (HNSCC), Colorectal cancer) and from different sequencing technologies (Smart-seq2, 10X Chromium), classifying a total of 106 samples and 93322 cells (Supplementary Table S2). In all the considered datasets, the identification of the non-malignant cell was reported by the authors and was based on a combination of approaches using copy number^4^, clustering and cell markers. We compare our results in terms of F1 score^20^ with those obtained by using a the state-of-the-art tool CopyKAT^11^. SCEVAN, as shown in figure 2, achieved a better classification score in 63% of the samples, whereas CopyKAT performed better than SCEVAN in 23% of the samples. The F1 score for all samples obtained with SCEVAN is 0.90 in contrast to the F1 score of 0.63 obtained with CopyKAT. SCEVAN showed a low F1 SCORE in samples with very few tumour cells (between 1 and 15), which are present mostly in the case of Head & Neck cancer dataset (Supplementary Table S2). For these cases, the separation of malignant and non-malignant cells exclusively based upon hierarchical clustering is notoriously challenging. This behavior is also a well-known limitation of CopyKAT^11^. Conversely, the greedy segmentation algorithm implemented in SCEVAN is particularly efficient. Indeed, a direct comparison of the execution times on the same data showed that SCEVAN is 2x to 7x faster (Supplementary Fig. 1). Collectively these results confirm that SCEVAN can accurately discriminate between tumor and normal cells in different solid tumors using the copy number profiles inferred from scRNA-seq. Moreover, SCEVAN is faster and more accurate than other state-of-the-art approaches.

**Figure 2.**
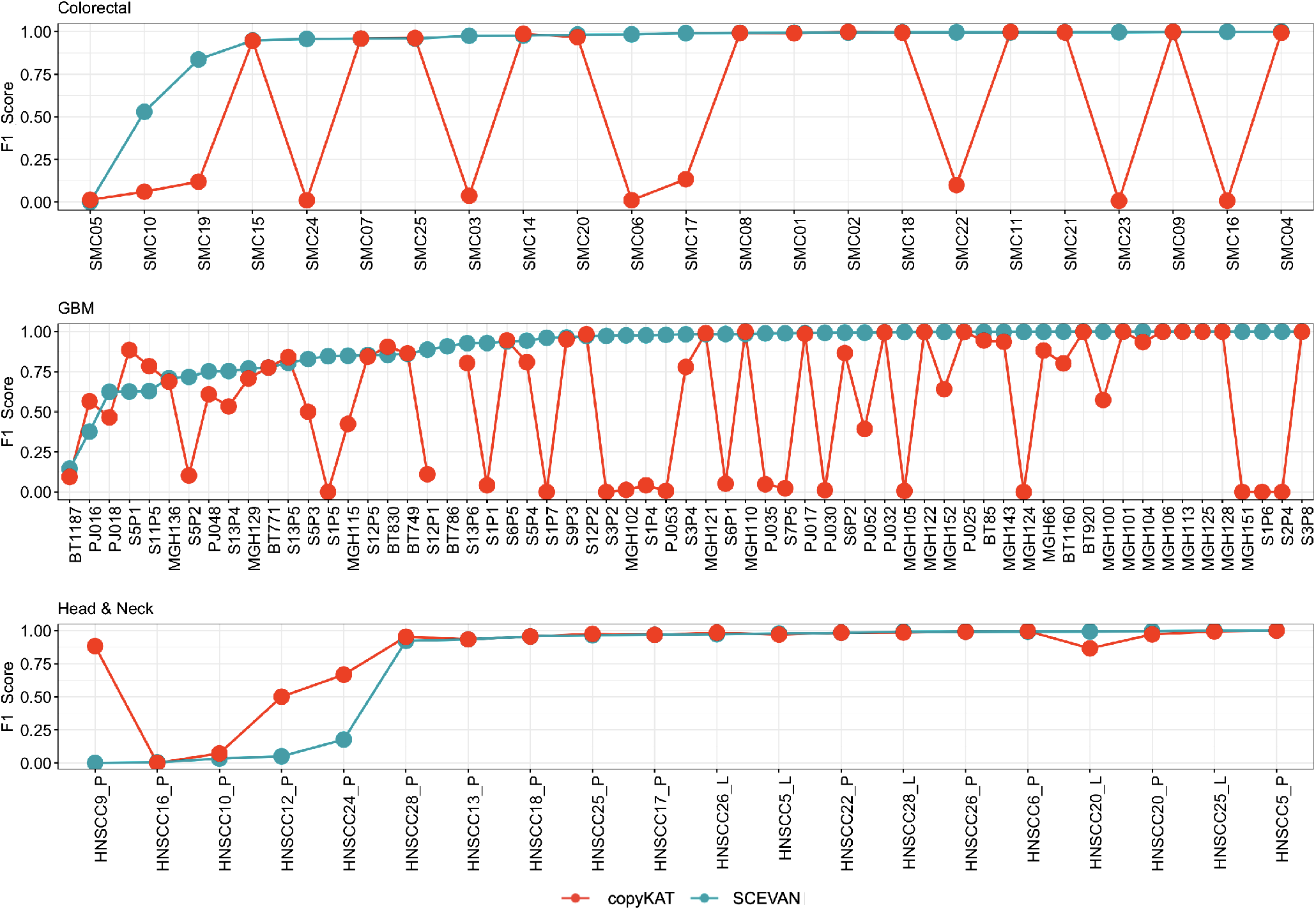
F1 score obtained with SCEVAN and copyKAT^11^ in the classification of malignant and non-malignant cells for each cancer type (Colorectal cancer^17^, Glioblastoma^7, 10, 19^, Head and Neck Squamous Cell Carcinomas^18^). Overall, 93, 322 cells from 106 samples (Supplementary Table S2).

### Accuracy of inferred copy number profile

We benchmarked the inferred copy number profile produced by SCEVAN on 25 samples of a Glioblastoma multiregional dataset^19^ against the CNV status obtained using low-depth whole-genome sequencing (WGS) on the bulk biopsies (Figure 3a). We re-sampled the output of SCEVAN and CopyKAT to the same resolution of the WGS by taking one value every 1 Mb. The Pearson correlation between the vector extracted from the segmentation and the smoothed ground truth reference was used for the evaluation. In the figure 3b the boxplots show the Pearson correlation obtained in all samples with the inferred profiles respectively with SCEVAN and Copy-KAT. SCEVAN has a mean correlation of 0.55 (max 0.73) whereas CopyKAT has a mean correlation of 0.06 (max). These data indicate that SCEVAN accurately infers DNA copy number profiles from high-throughput scRNA-seq data.

**Figure 3.**
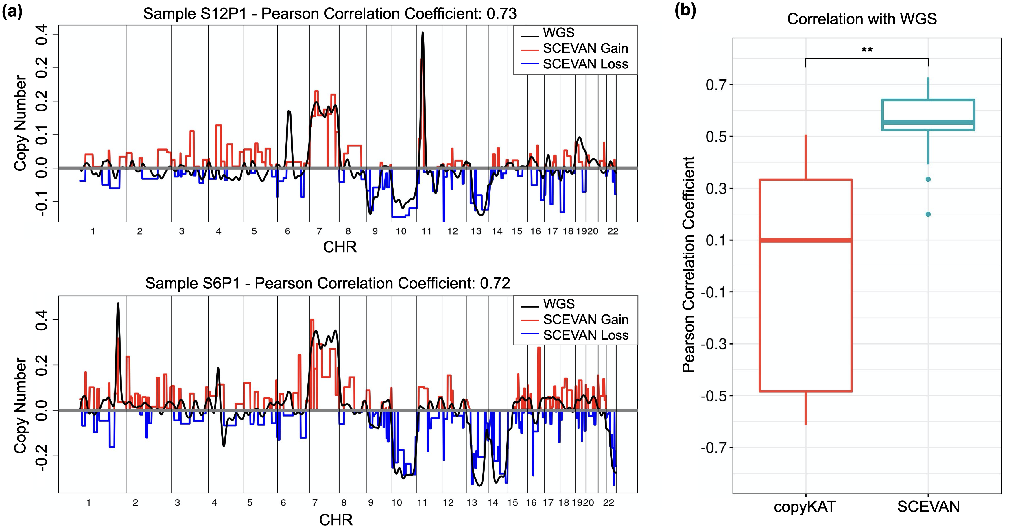
**(a)** Copy number profile inferred with SCEVAN and the corresponding ground truth from WGS of samples S12P1 and S6P1^19^. **(b)** Pearson correlation between the copy number inferred with SCEVAN or with CopyKAT and the ground truth from WGS for 25 samples^19^.

### Intratumoral heterogeneity in Glioblastoma

Glioblastoma (GBM) is the most aggressive form of brain tumor. The presence of clonal and sub-clonal differentiated tumor cell populations, glioma stem cells, and components of an immuno-repressive tumor microenvironment hinders the definition of effective therapies^21^, such heterogeneity is influenced by multiple genetic and epigenetic clues^7, 22^. SCEVAN can automatically infer clonal substructure from single-cell data by analyzing the clusters of the CNA matrix that show significantly different genomic alterations (Methods). As an application of this approach, we considered one of the samples reported in a recent study^7^, the MGH105 sample. SCEVAN identifies four subpopulations that have different alterations on chromosome 6 (Supplementary Fig. 2). Interestingly, whereas the resolution for the identification of four subclones could not be reached by canonical scRNA processing analyses^7^, the existence of the same subclones detected by SCEVAN using exclusively single cell transcriptomic data in this tumor was recently confirmed by the application of DNA single cell DNA methylation platforms^22^. Taken together, these findings highlight the superior performance of SCEVAN that shows an unprecented resolution exclusively from scRNA-seq datasets up to a level that was previously obtained only through concurrent application of multi-omics single cell data.

In yhe sample BT1160, the execution of SCEVAN returns the presence of three tumor cell sub-populations, as shown in figure 4a-b. Phylogenic reconstruction of the clone tree shows that two of the clones are similar (subclone 1 and 2) whereas the third subclone is significantly far (Figure 4c), suggesting different dynamics of clonal expansion and diversification. In order to better understand how individual clones fuel tumor growth and clonal selection, we investigated the reported alterations. SCEVAN identifies several truncal alterations, such as the amplification on Chr 5 (q23.2-q31.3), shared alterations, such as the deletion on Chr 10 (q22.1-q26.3), and subclone-specific alteration such as the amplification in the green sub-population on Chr 1 (q31.2-q32.1) and Chr 19 (q13.32-q13.33) ((Figure 4d). Interestingly, subclone-specific functional analysis reveals a differential activation of pathways that resemble a recent metabolic classification of Glioblastoma^5^. Subclone 1 (lightblue) enriches pathways characteristic of the Neuronal subtype, subclone 2 (blue) has cells belonging to the Mitochondrial, and the subclone 3 (green) contains cells with Proliferative/Progenitor subtype (Figure 4e). Indeed, this finding is also confirmed by the enrichment of individual cells for every subtype (Figure 4f). The evolutionary reconstruction suggests the hypothesis that the subclone 3 is the funding clone. The Proliferative/Progenitor subclone has several specific amplifications (1q21.3-q22, 1q31.2-1q32.1, 3q26.32-3q27.2, 4q32.1-4q35.1, 6p22.1, 8p11.22-8q21, 19q13.32-19q13.22). To identify drivers of the different cellular states we performed differential analysis between genes with genomic coordinates in regions of the subclone-specific alterations. The top differentially expressed gene lying in the alterations specific of the subclone 3 was Ubiquitin-conjugating enzyme E2T (UBE2T) gene, which is significantly up-regulated (p-value 2.69*e*^−43^ log fold change 1.10) (Supplementary Fig. 3) enriching the activity of the pathway of DNA Repair. This gene encodes for the exclusive ubiquitin-conjugating enzyme (E2) that partners with the Fanconi Anemia (FA) ubiquitin ligase (E3). The E2T-FA complex is required for DNA interstrand crosslink repair as the monoubiquitination event implemented by E2T is essential for the recruitment of downstream DNA repair factors by FA^23^.

**Figure 4.**
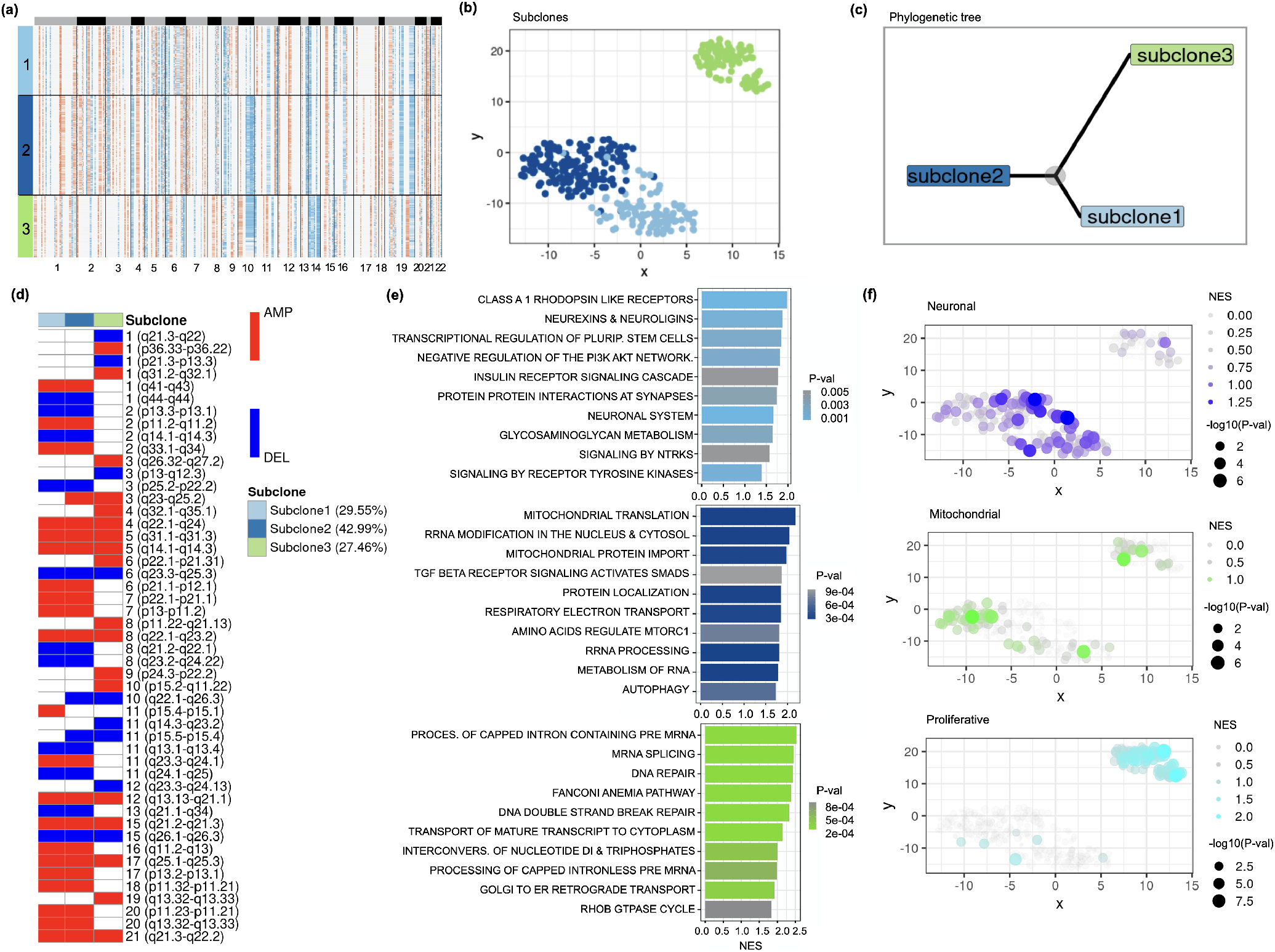
**(a)** Clonal structure of sample BT1160 inferred by SCEVAN. **(b)** t-SNE plot of CNA matrix. **(c)** Inferred phylogenetic tree. **(d)** OncoPrint-like plot of BT1160 highlighting clone-specific alterations, shared alterations between and clonal alterations. **(e)** REACTOME pathways activity for each subclone. **(f)** NES per cell of GBM cellular states^5^.

Furthermore, the analysis of copy number sub-structure can characterize the clonal status of specific tumor-associated genes. In particular, SCEVAN reveals that in samples BT1160 and MGH102, alterations of tumor suppressor genes CDKN2A and PTEN are subclonal (Figure 5). Indeed, in sample BT1160, the deletion on Chr 10 (q22.1-q26.3), containing PTEN (10q23.31), is shared between two out of three subclones, while in the remaining subpopulation, this alteration is not present. Also, in the sample MGH102, the region 9p21.3 containing the gene CDKN2A is deleted in two of the four subclones. Taken together, these results suggest that SCE-VAN can resolve clonal copy number substructure in tumors from scRNA-seq data and identify subclonal differences and glioma-specific cancer states.

**Figure 5.**
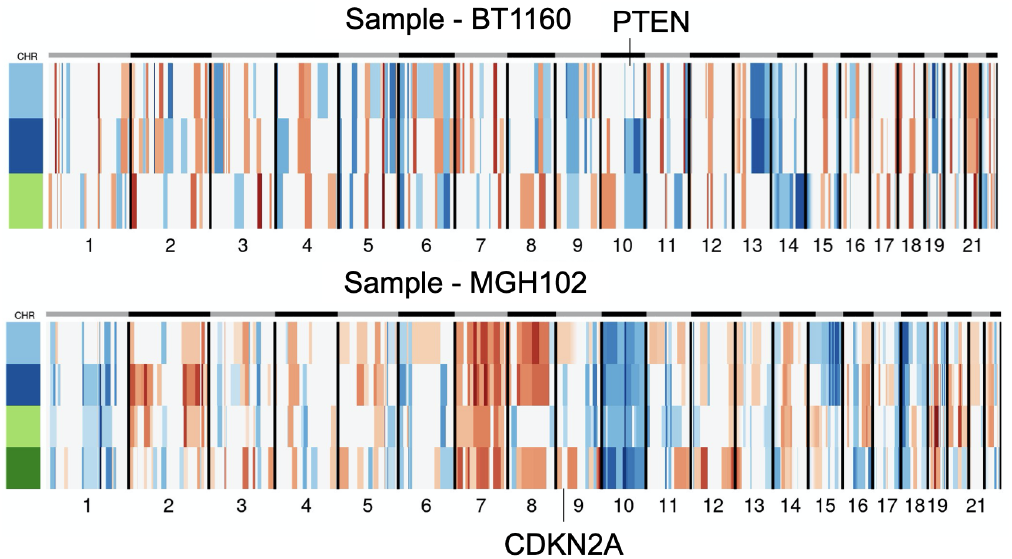
Compact representation of clonal structure inferred with SCEVAN of scRNA samples BT1160 and MGH102^7^, in which the alterations containing tumor suppressor genes PTEN and CDKN2A are subclonal.

### Clonal evolution in multiregional GBM tumor

Glioblastoma heterogeneity has also been investigated in the spatial and temporal axes^19, 24^ because a single biopsy may not be informative of the whole tumor. Multiple biopsies allow us to characterize the clonal architecture and evolutionary dynamics of GBM^25^.

We use SCEVAN for evolutionary analysis of clonal structure for multi-regional scRNA-seq samples of GBM^19^. For example, we consider one case, GS1, with seven biopsies, two of them taken at the tumor periphery and the remaining at the core of the tumor. The clonal analysis of each sample with SCEVAN allows inferring an evolutionary tree of the clones (figure 6). Copy number alterations develop along several branches, and the peritumoral samples (P2/P3) are in a branch separated from the core samples, in which there is no amplification in chromosomes 4 and 8. Moreover, the amplification present on Chr 2 is clonal in peripheral samples and is subclonal in some core samples (P1/P4/P7).

**Figure 6.**
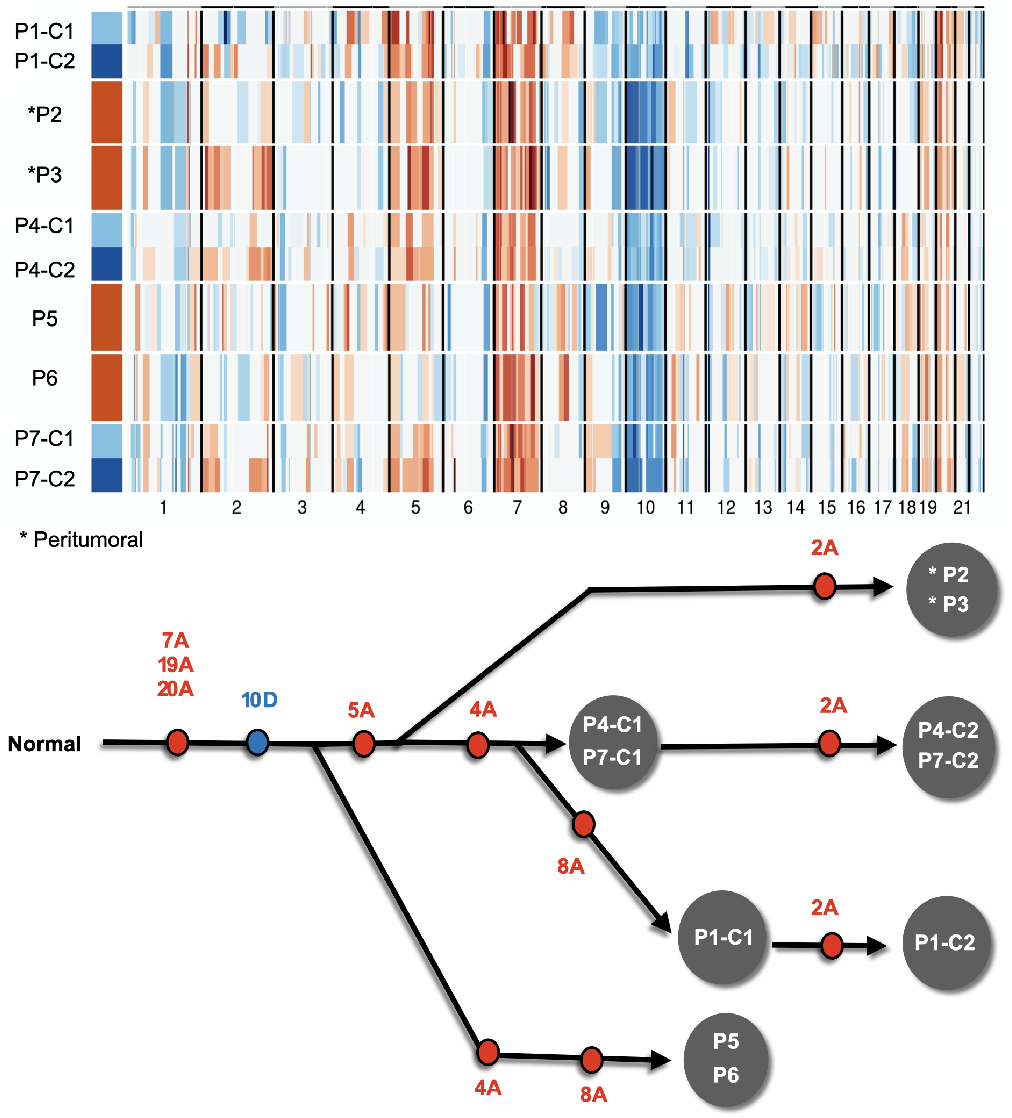
Compact representation of clonal structure inferred with SCEVAN of multi-regional scRNA-seq samples of patient GS1^19^ and a phylogenetic tree deduced from clonal structure of the samples.

### Clonal structure of primary and metastatic lymph

SCEVAN (and similar approaches) can address important questions such as identifying similarities and differences between primary tumors and metastases. For this purpose, we consider primary HNSCC tumors and corresponding lymph node metastases^18^. Of the four considered cases, just one specific sample, patient (HNSCC5), presents a different clonal structure between primary tumor and lymph node metastasis, in particular the absence of amplification of chromosome 7 (p22.3-p13) in the lymph node metastasis, as shown in figure 7. Interestingly, this is the locus of Glycoprotein non-metastatic b (GPNMB) which is down-regulated in the lymph node metastasis (Supplementary Fig. 4). Furthermore, GPNMB increases tumor growth and metastasis in multiple contexts^26^. For the remaining patients (HNSCC20,HNSCC25,HNSCC26,HNSCC28) the clonal structure of the lymph node metastasis appears to be the same as in the primary tumor. Therefore, we obtain a high correlation (Pearson correlation between 0.79 and 0.89) comparing the clonal profiles of the primary tumor and lymph node metastasis pairs. These data show that result obtained with SCEVAN can be used to study the clonal evolution of metastatic cancer.

**Figure 7.**
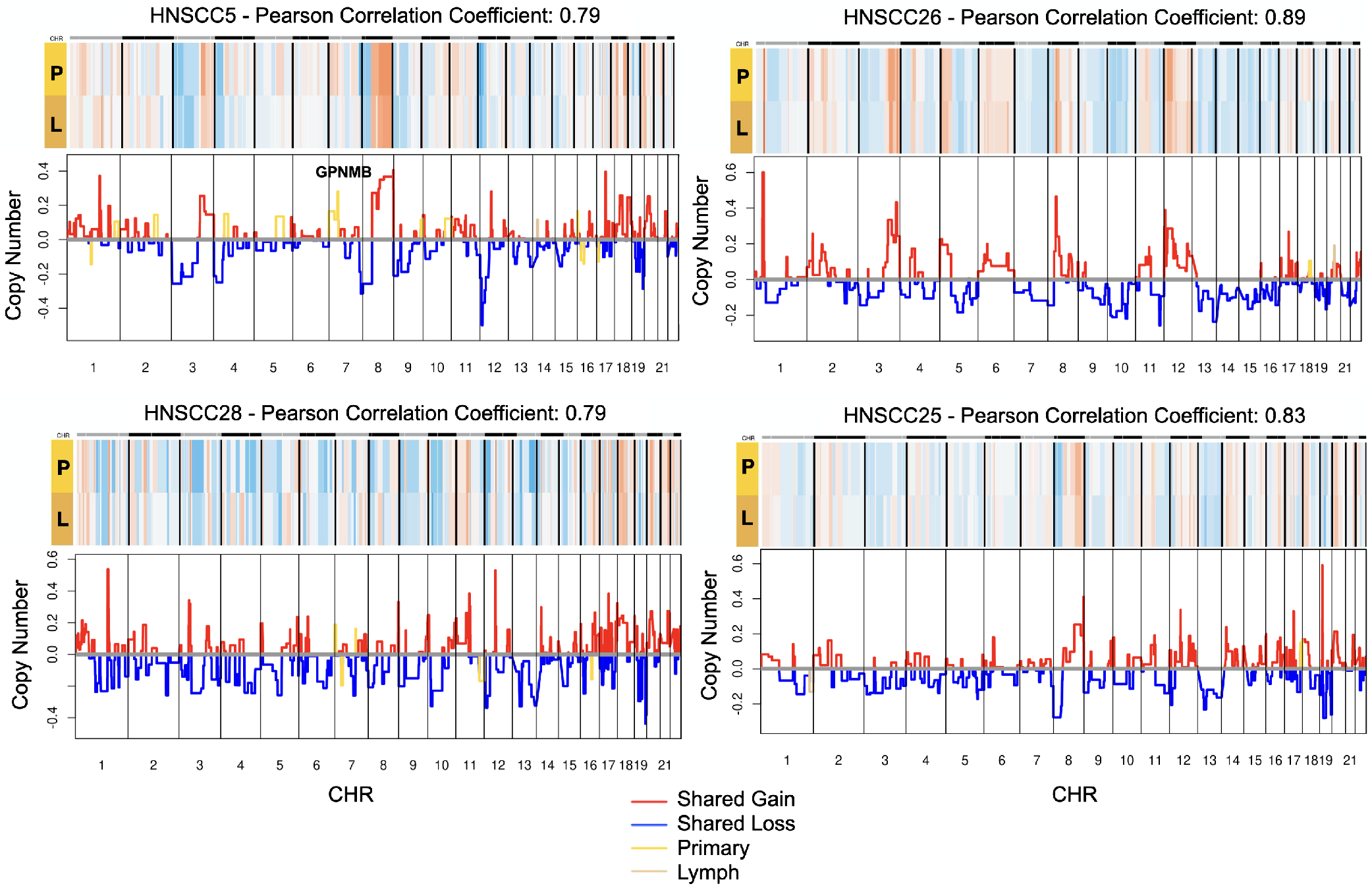
Copy number profile of Primary (P) and Metastatic Lymph nodes (L) from samples of Head and Neck cancer dataset (HNSCC5,HNSCC25,HNSCC26,HNSCC28)^18^.

## Discussion

We described a variational segmentation approach to identify genomic copy number profiles from scRNA-seq data. The adopted joint segmentation algorithm is based on the notion that the cells in a given copy number clone share the same breakpoints. Thus, the expression profile of every individual cell, seen as a function of the genomic coordinates, contributes to the evidence of the presence of copy number alteration in each subclone. SCEVAN uses a set of stromal and immune signatures and the fact that malignant cells often harbor aneuploid copy number events to discriminate between transformed cells and micro-environment cells automatically. We used an extensive collection of annotated datasets of different tumor types confirming that SCEVAN is more accurate and faster than state-of-the-art methods. Our evaluation has shown that this approach is viable in cases with very high purity and subjects with a significant amount of immune infiltration. Therefore, SCEVAN is particularly suited in studies where unsorted populations of single cells need to be analyzed to characterize, for example, the interaction between malignant cells and their microenvironment^6^.

The primary use of SCEVAN consists of delineating the clonal substructure in solid tumors based on differences in CNAs and studying the temporal and geographic evolution of tumors. In addition, we used SCEVAN to deconvolve the clonal structure of glioma tumors. For example, in one patient, we found the presence of cell populations with differential activation of glioma cellular states, confirming that the heterogeneity of glioma subtypes is driven by the clonal architectures^5^. Functional analysis of subclones revealed novel drivers of cellular states such as the Proliferative/Progenitor (PPR) glioma subtype. We identified UBE2T as the top amplified and differential expressed gene in the PPR clone. Interestingly, UBE2T can be pharmacologically inhibited^27^, and therefore it results as a new potential therapeutic target of PPR cells. Moreover, we have shown that with SCEVAN, we can characterize the clonal status of onco-suppressor genes such as PTEN and CDKN2A. Such characterization may be of interest for diagnostics or therapeutic targeting and for exploitation of approaches based on synthetic lethality^28^. Clonal deconvolution extracted from scRNA-seq can also be used to study regional and temporal tumor evolution as we have shown in the case of a multiregional GBM dataset and for the characterization of difference between primary and metastases.

Some limitations of our SCEVAN rely mainly on its basic assumption that their aneuploidy can identify cancer cells. However, there are cases such as liquid cancers ((es. Leukemia), pediatric cancers, Ependymomas, and others are known to harbor a minimal number of genomic alterations. Thus, our approach (and similar) may not be suited in this case.

## Methods

### Preprocessing of scRNA-seq data

The preprocessing phase is aimed at filtering out low-quality and irrelevant cells. Specifically, the cells with less than 200 detected genes and the genes expressed in less than 1% of cells are removed. The remaining genes are annotated by adding their genomic locations to the matrix using Ensembl based annotation package^29^ and then genes are sorted according to genomic coordinates. After annotation, the genes involved in the cell cycle pathway, obtained from REACTOME^30^, are filtered to reduce artificial segments caused by the cell cycle^11^.

### Identification of High confident non-malignant cells

To segregate malignant from non-malignant cells, SCEVAN follows a multi-step approach. A small set of high confidence normal cells is used to build a relative expression matrix and seed the cluster of other normal cells. Then, the relative expression matrix is segmented and then clustered as described in the following paragraphs. To identify the high confidence normal cells, we use a set of gene signatures from public collections^6, 31^, which includes cells of tumor microenvironment including stromal and immune cells, such as lymphocytes, macrophages, microglial cells, dendritic cells, neurons, and others (Supplementary Table 1). We apply the Mann-Whitney-Wilcoxon gene set test gene set enrichment analysis implemented in the yaGST package^32^ and assume as normal confident cells the top classified cells with *p*-value less than 10^−10^ and Normalized Enrichment Score greater than 1.0. We restrict the search to a maximum of 30 high confidence non-malignant cells. Then the copy number baseline, estimated from the median expression of confident normal cells, is removed from the count matrix, thus obtaining the relative matrix 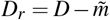 where 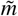 is the baseline *n*-dimensional vector (*n* being the number of genes) with the median value of confident normal cells.

### Edge-preserving smoothing

Before the segmentation phase, one of the key steps of SCEVAN is to smooth the relative expression matrix. Since the segmentation step described below assumes a piecewise-constant model of the copy number signal, we preliminarily proceed to perform a nonlinear smoothing of the gene expression along with the genomic coordinates to regularize the signal, reduce the outliers and at the same time preserve the discontinuities which are the breakpoints between the copy number segments. We apply a filter grounded in the Bayesian framework of edge-preserving regularization^33^ which considers the minimization of the Total Variation (TV) functional

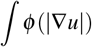

where *u* is the relative gene expression signal, ∇*u* is its gradient and *ϕ*(·) is a discontinuity-adaptive prior^34^. In particular, here we use *ϕ*(*x*) = logcosh(*x*) which has been shown to produce a well-posed minimization problem overcoming the non-differentiability of the TV at the origin^35^. The iterative numerical scheme implemented in SCEVAN is just the one-dimensional adaptation of the stable finite difference scheme previously reported^35^.

### Single Cell joint segmentation algorithm

SCEVAN uses a joint segmentation procedure that inputs all the cells in a given clone to identify the boundaries of homogeneous copy number. Standard segmentation procedures independently segment each sample^11^. The procedure is based on the *Mumford and Shah energy* originally developed to analyze images. In their original work^12^, the authors introduced the basic properties of variational models for computer vision aimed at defining the mathematical foundations for appropriately decomposition of the 2D domain Ω of a vector-valued function *u*_0_ : Ω → ℝ^*m*^ into a set of disjoint connected components (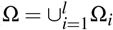, Ω_*i*_ ⋂ Ω_*j*_ = ∅, 1 ≤ *i,j* ≤ *l, i* ≠ *j*). The set of points on the boundary between the Ω_*i*_ is denoted as Γ. This partition is modeled such that the signal varies smoothly within a component and discontinuously between the disjoint components. This problem is known as piece-wise smooth approximation. For this purpose, Mumford and Shah proposed the minimization of the following functional:

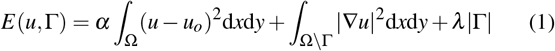

where *α* and *λ* are two non negative parameters weighting the different terms in the energy: the first term requires that *u* approximates *u*_0_, the second term takes in account the variability of *u* within each connected component Ω_*i*_ and the third term penalizes complex solutions in terms of the length of the boundaries |Γ|. Here we adopt a special case of equation (1) when the approximation *u* of the signal *u*_0_ is constrained to be a piece-wise continuous function (*u* constant within each connected components Ω*i*). This is best suited for CNV segmentation. In this case, the second term of the energy functional vanishes, so the optimal segmentation is obtained by minimizing:

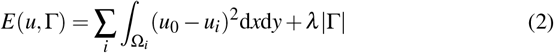

in this case, it easy to show that the minimum for this model can be obtained by posing *u_i_* as the mean of *u*_0_ within of each connected component Ω*i*. Hence, this functional represents a compromise between the accuracy of the approximation and parsimony of the boundaries. It is essential to notice that the resulting segmentation depends on the scale parameter *λ*. Indeed, it determines the number of computed regions: when *λ* is many small boundaries are allowed, the resulting segmentation will be fine. As *λ* increases, the segmentation will be coarser and coarser. Specifically, the input data *D* is an *m* × *n* single-cell gene expression matrix obtained after the preprocessing step detailed above, where *m* is the number of cells in a given clone, and *n* is the number of genes ordered by genomic positions. Then, we define a segmentation *D* as a set of ordered positions (breakpoints) partitioning the columns of D into *M* connected regions *R* = {*R*_1_, …, *R*_*M*_} identified by a set of ordered positions Γ = {*b*_1_,…, *b*_*M*+1_}. Each region *R*_*i*_ contains all genes whose genomic coordinates lie between breakpoints {*bi, b*_*i*+1_}. Moreover, we are dealing with the one-dimensional vector function where the domain is the genome, therefore in equation (2) |Γ| reduces to the number of regions *M*. According to the original algorithm proposed in^16^, to minimize this function, adjacent regions *R*_*i*_ and *R*_*i*+1_ are iteratively merged in a pyramidal manner to create larger segments, and the reduction of the energy can be shown as:

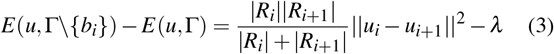

where *R*_*i*_ is the lenght of the i-th region, and *u*_*i*_ is an *m*-dimentional vector with the mean value of these columns, || · || is the *L*_2_ norm and is \ the set difference. To minimize (2), we follow a greedy procedure. We start with a segmentation having *n* regions, one for each gene. Then, at each step, we merge the pair of adjacent regions that yield the maximum decrease of the energy functional upon merging. Since *λ* decides the end of merging, the choice of an appropriate value is crucial to ensure the quality of the final segmentation. As in^16^, the selection for *λ* at each merging step is done dynamically, depending on two factors - the size of the region and mean values of the consecutive regions being considered for the merge. Hence, the cost of merging two regions *R*_*i*_ and *R*_*i*+1_, associated with a breakpoint *b*_*i*_, is computed as follows:

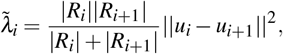

if 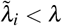 the adjacent regions are merged and the i-th breakpoint removed. Otherwise, the energy function has reached a local minimum, and no merging can be done further. Therefore, *λ* is updated to the smallest of *λ*_*i*_ + *ε*, continuing the merging. The sequence of *λ* values is monotonically increasing as it corresponds to the amount of decrease of the energy functional at each step in (eq. (3)). In^13^ we adopted a stopping criterion in such a way that the final segmentation is obtained when the increase in *λ* stabilizes and merging any further does not correspond to a significant decrease of the energy. The final stopping value is based on the variability of the adjacent region and the total variability of the data, *v*. The resulting computation for the stopping criterion is Δ*λ* = *λ*_*i*+1_ − *λ*_*i*_ ≤ *β v*, where *β* is a positive constant (ideal value = 0.5–0.7), which represents the only parameter of the segmentation algorithm.

Finally, the joint segmentation procedure performs a significance test that evaluates the statistical significance of the alterations (amplification o deletion) observed in a given region. The null hypothesis of the test is that the *i*-th region has no CNA, expressly assuming *μ* as the number of cells with the alteration in the region and *μ*_0_ as the expected count when no alteration is present in a region, we have *ℋ*_0_ : *μ* = *μ*_0_ and *ℋ*_1_ : *μ* > *μ*_0_. Indeed the null distribution is estimated from the binary matrices (*N* cells and *M* segmented regions), in which the value 1 indicates that a cell shows an alteration in the *i*-th region if the mean is above a fixed threshold. Then the distribution of the test statistics is computed via random permutations as previously described^15, 16^.

### Classification of malignant and non malignant cells

The multi-valued function *u* resulting from the segmentation is computed by substituting each value with the mean between consecutive breakpoints in each cell. It is then clustered into two groups using hierarchical clustering. All the cells in the cluster containing the highest number of confident normal cells are then classified as non-malignant. The final CNA matrix is then obtained subtracting the mean value of all the identified normal cells from each genomic position.

If no confident normal cells are found, we assume that the sample is pure and contains only malignant cells. In this case, the CNA matrix is obtained by removing from the malignant cells a synthetic baseline.

### Differential sub-clonal structure characterization

SCEVAN can automatically characterize sub-clonal structures in tumor cells. Cancer cells are clustered according to their copy number profile. We use Louvain clustering^36^ applied to a shared nearest-neighbor graph^37^ of the first 30 principal components of the final CNA matrix. Each cluster represents a potential subclone. Therefore the joint segmentation algorithm is re-applied considering just the cells of the cluster, the segmentation results are analyzed to identify subclone specific alterations, shared alterations between subclones, and clonal alterations. From the segmentation of each subclone, we select segments representing relevant alterations with |*u*| ≥ 0.10 or significance testing of the joint segmentation with *p*-value ≤ 0.05 and then a hierarchical aggregation is carried out in which the neighboring segments are aggregated together if they are of the same kind (amplifications or deletions) and if their genomic distance is less than 10*Mb*. This hierarchical aggregation serves to achieve a coarse-grained segmentation between subclones. The aggregated alterations extracted for each subclone are compared among subclones to determine specific or shared alterations. For each subclone, we perform a linear search for alterations in the other subclones. An alteration is shared between if the respective start or end breakpoints are at a genomic distance of less than 10*Mb* and if the length of the minor segment is at least 40% of the size of the major segment.

The potential specific alterations and those shared only between certain subclones, obtained previously, are further investigated by performing a significance test between the corresponding segments among the clones. Each alteration is considered specific if the *p*-value (t-test) between the mean of the genes in the CNA array, belonging to the segment, of the cells presenting the alteration and those not presenting the alteration is less than 10^−10^ and the difference in the mean is greater than 0.05.

## Code and Data availability

SCEVAN is available in open source as an R package at the following address https://github.com/AntonioDeFalco/SCEVAN.

Data used in this paper are publicly available on the Gene Expression Omnibus (GEO) with accession numbers listed in the Supplementary Table 2. Copy number data from low pass whole-genome sequencing are available contacting the authors of^19^.

---

**Figure S1.**
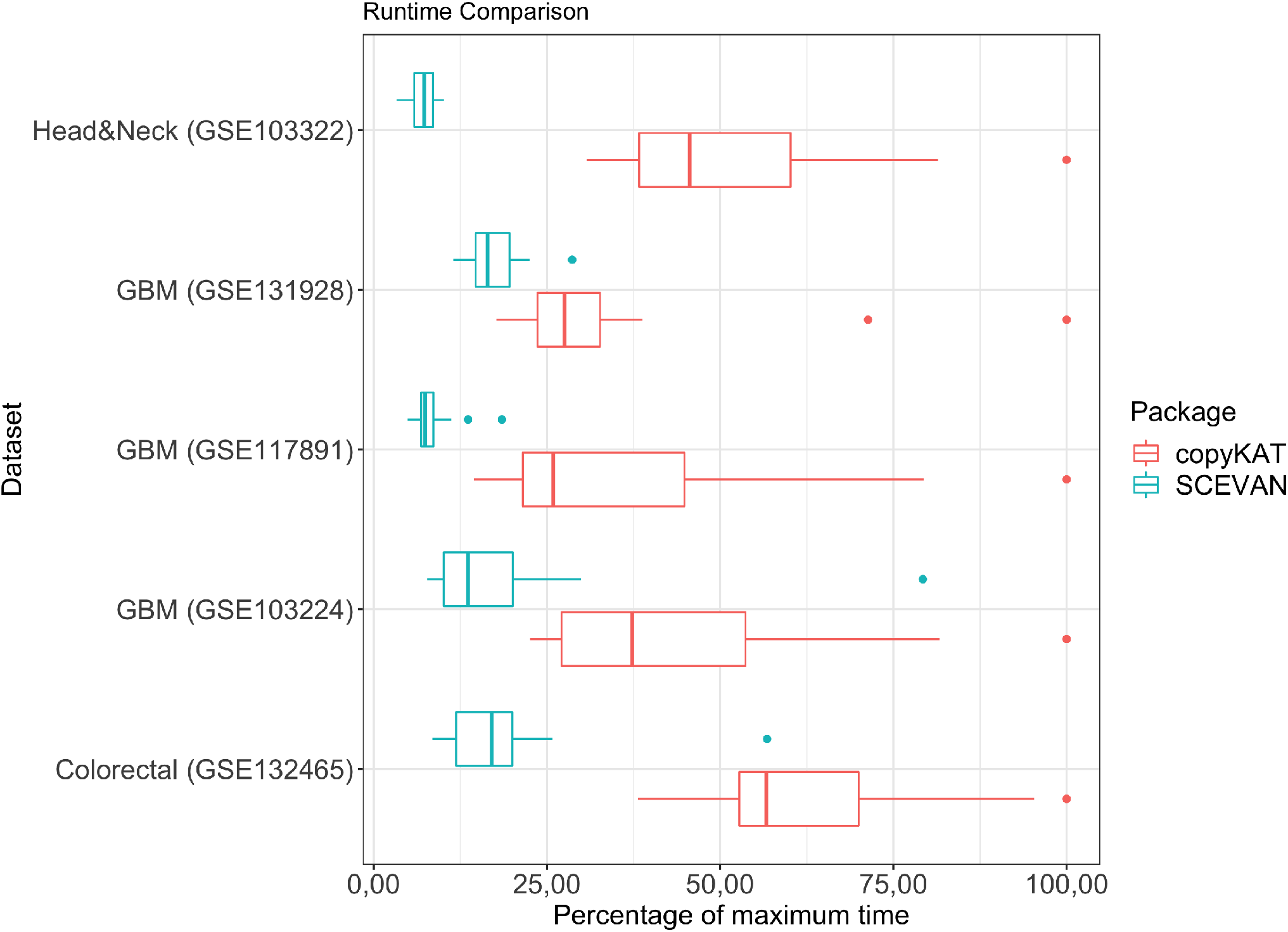
Classification time for each sample expressed as a percentage relative to the maximum time for each dataset.

**Figure S2.**
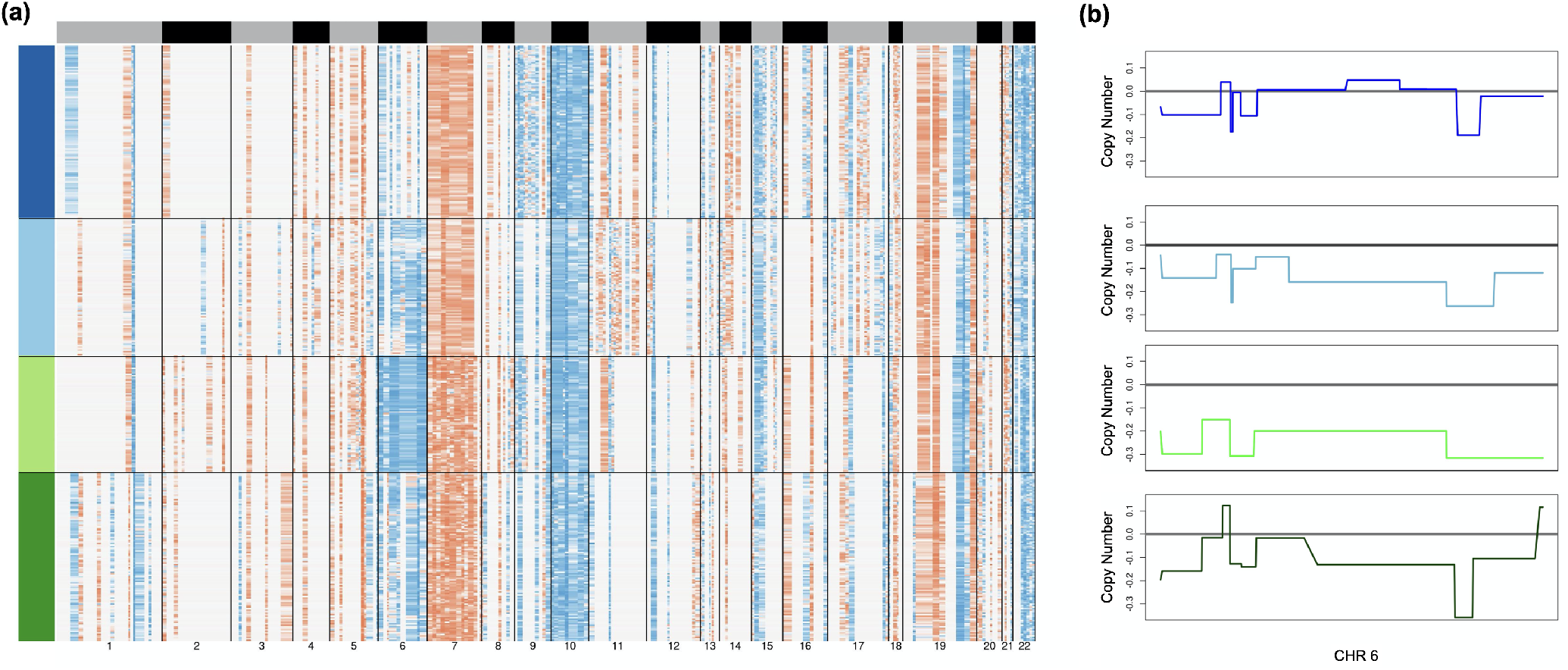
**(a)** Clonal structure of MGH105^7^ inferred by clustering single-cell copy number profiles by SCEVAN. **(b)** Inferred copy number in chromosome 6 for each subclone.

**Figure S3.**
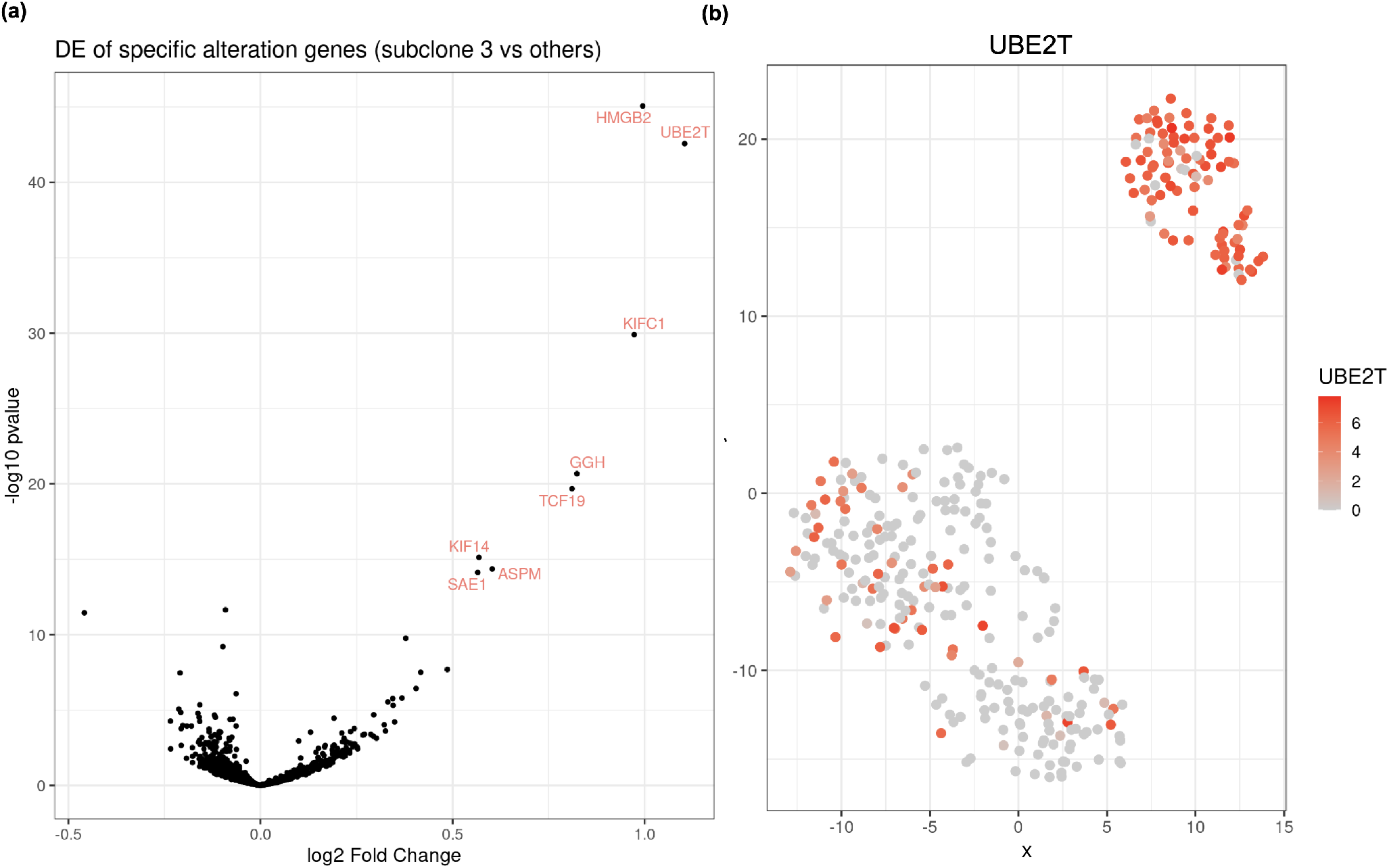
**(a)** Differential gene expression analysis of genes belonging to the specific amplifications of subclone 3, comparing subclone 3 against the others. **(b)** UBE2T expression on t-SNE plot of CNA matrix.

**Figure S4.**
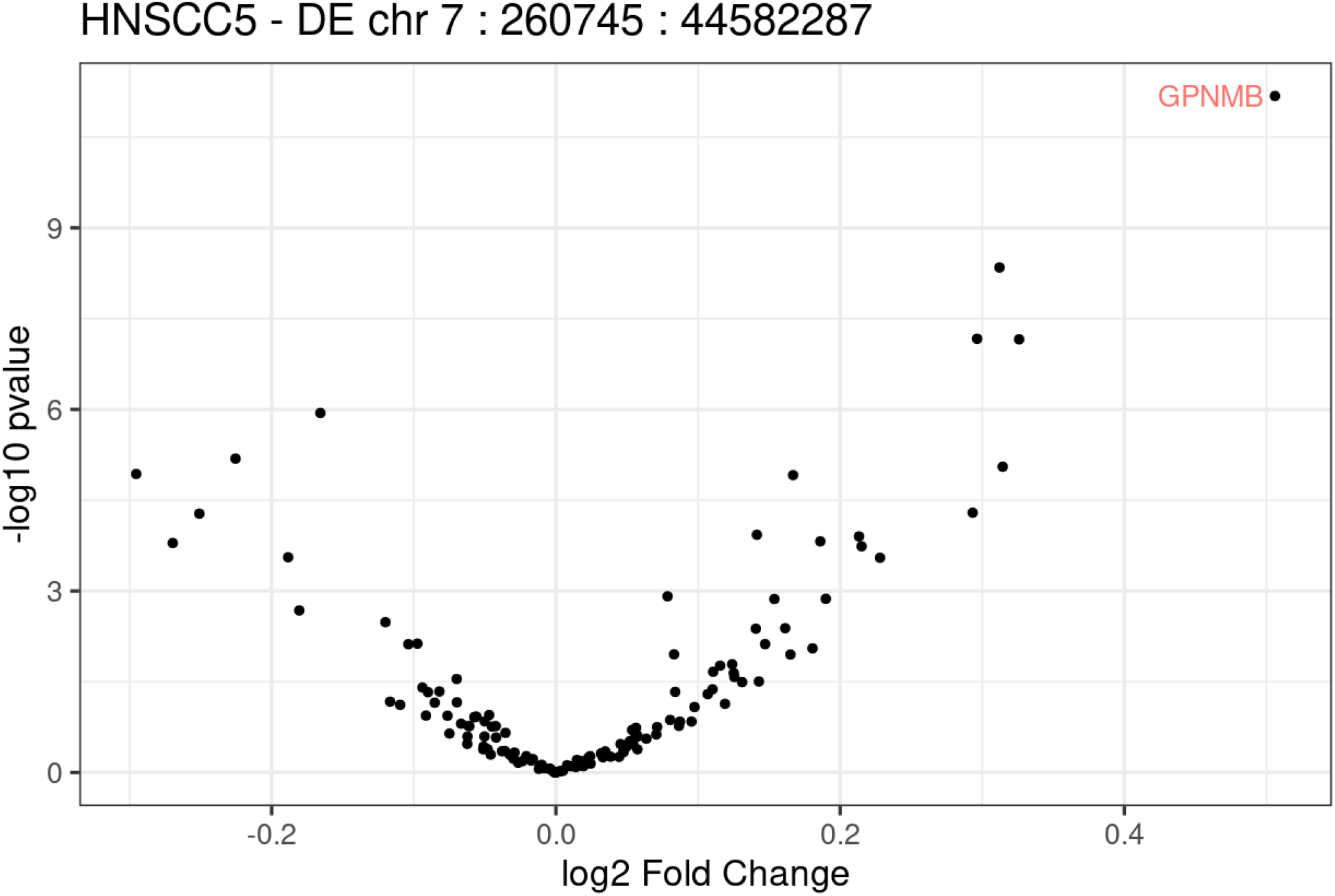
Differential analysis of genes belonging to the specific amplification on chromosome 7, between the gene expression of primary tumor against the lymph node metastasis.

